# Precise Functional Localization of the Foveolar Representation in Anesthetized Macaque Visual Cortex

**DOI:** 10.64898/2026.08.16.744304

**Authors:** Meizhen Qian, Meixuan Chen, Annamaria Barczak, Xiaotong Zhang, Peichao Li, Anna Wang Roe

**Affiliations:** Department of Anesthesiology of the Second Affiliated Hospital & Liangzhu Laboratory, Zhejiang University School of Medicine, Hangzhou, China; Interdisciplinary Institute of Neuroscience and Technology, Zhejiang University School of Medicine, Hangzhou, China; Division of Translational Neuroscience, Center for Biomedical Imaging and Neuromodulation (C-BIN), Nathan S. Kline Institute for Psychiatric Research, Orangeburg, NY; MOE Frontier Science Center for Brain Science & Brain-Machine Integration, State Key Laboratory of Brain-machine Intelligence, School of Brain Science and Brain Medicine, Zhejiang University, Hangzhou, China; NHC and CAMS Key Laboratory of Medical Neurobiology, Zhejiang University, Hangzhou, China; College of Electrical Engineering, Zhejiang University, Hangzhou, China; Departments of Psychiatry and Neuroscience, NYU Grossman School of Medicine, New York, NY

**Author notes:** Co-first authors: These authors contributed equally (Meizhen Qian, Meixuan Chen). Corresponding Authors: These authors jointly supervised this work, Anna Wang Roe, Director of Translational Neuroscience Laboratory Division, Center for Biomedical Imaging and Neuromodulation (C-BIN), Nathan S. Kline Institute for Psychiatric Research, Meizhen Qian, Research Scientist, Translational Neuroscience Laboratory Division, Center for Biomedical Imaging and Neuromodulation (C-BIN), Nathan S. Kline Institute for Psychiatric Research, Peichao Li, Professor, Interdisciplinary Institute of Neuroscience and Technology, School of Medicine, Zhejiang University, Hangzhou, China.

## Abstract

The foveola—the central ∼1° of the visual field—supports the highest-acuity spatial vision, yet localizing its cortical representation is technically demanding, particularly under anesthesia where gaze cannot be behaviorally controlled. Here we describe a hierarchical, meridian-based 7T BOLD fMRI strategy for sub-degree localization of the foveolar representation in anesthetized macaques. Drifting grating bars presented at three successively finer scales (4.5°, 1.5°, 0.5°) independently refined the vertical and horizontal meridian representations, each identified by a characteristic activation signature: bilateral convergence at the V1/V2 border for the vertical meridian, and simultaneous dorsal and ventral V2/V3 activation at the fundus of the calcarine sulcus for the horizontal. Their functional intersection defined candidate foveolar coordinates, which were refined using 0.2° spot stimuli. Surface-based rendering resolved discrete foveolar loci at the lateral end of the V1/V2, V2/V3, V3/V4 and V4/TEO borders, together delimiting the previously described ‘foveolar core’. Pooling these with awake data, we place the loci in a common stereotaxic frame, yielding a first anatomical atlas of the core. Intrinsic-signal optical imaging further shows that this territory lacks the ocular-dominance and orientation columns of surrounding cortex. The approach provides a reproducible framework for locating and targeting foveolar cortex in anesthetized primates.

## Introduction

The primate visual system devotes a disproportionately large amount of cortical territory to processing information from the central visual field^1–3^. Within this region, the *foveola*— representing the central 1° of visual angle—plays a fundamental role in high acuity spatial vision, color discrimination, and gaze-dependent attentional selection^4–6^. The unique anatomical and functional properties of the foveola, including the highest cone density in the retina and strong cortical magnification in early visual areas^2,6,7^, enable primates to perform visually guided behaviors such as object recognition, reading, and fine motor coordination^4,8^. Understanding how the cortex represents and processes foveolar inputs is therefore essential for elucidating the neural mechanisms underlying central vision and visual cognition^9,10^.

Precise functional localization of the visual foveola in the primate cortex remains technically challenging. Retinotopic mapping techniques have been extensively used to delineate visual area boundaries^11–14^. These methods are typically optimized for identifying large-scale topographic organization rather than achieving sub-degree precision at the foveal center. In awake, behaving monkeys, foveolar localization can be facilitated by fixation control and eye tracking, but stable behavioral fixation requires extensive training^15^. In contrast, anesthetized preparations can provide physiologically stable conditions, without signal artifacts due to body motion, making it suitable for high-resolution imaging^16–18^. However, unlike in the awake animal, in which both eyes are directed to the location of gaze, the gaze direction of the eyes, which can differ substantially for the left and right eyes, is challenging to accurately determine under anesthesia. Placing visual stimuli on a tangent screen or monitor for studying foveolar cortical responses then becomes potentially fraught with error. Typically, the position of foveolar gaze on the tangent screen can be approximated by back-reflecting either the foveal center (which appears dark on the retina due to photoreceptor density) or the optic disk and then measuring the distance to the foveal center (14° nasal and 1° inferior) on the tangent screen; however, this approach provides at best only a coarse (±1-2° of error) estimate of central gaze^19^. In large eyes, such as that of cats, the vascular pattern can be reflected onto the tangent screen^20^; however, in monkeys this is difficult because the foveal avascular zone leaves no vascular landmark at the foveal center and the sub-degree precision required to localize the foveola exceeds that achievable by vascular back-projection. Thus, a more precise gaze mapping method under anesthesia is needed.

An additional and often underappreciated difficulty arises from the anatomical location of the foveolar cortex itself. In 1969, Zeki predicted the presence of a ‘foveal confluence’^21^, a term referring to the foveal juncture of V1, V2, V3, and V4. This confluence has subsequently been studied using neuroimaging methods in humans and monkeys, revealing either a single region of foveal representation shared by V1-V4 or multiple foveal representations, one locus for each early visual area ^22,23^. More recently, using ultrahigh field MRI imaging in awake, fixating monkeys, we found the confluence comprises 8 foveolar loci per hemisphere (one each at the V1/V2, V2/V3, V3/V4, and V4/TEO borders, both dorsally and ventrally), and bounded by these loci a region we term the ‘foveolar core’. While it is unknown whether a similar arrangement is present in human foveolar cortex, careful perusal of previous neuroimaging studies hints at a similar presence of multiple foveolar representations at the foveal confluence^24,25^, a possibility that requires further study. This relatively large foveolar complex in the macaque monkey is positioned at the extreme lateral pole of the occipital operculum^26^, a location that can make it challenging for electrode access^27^; and even when accessed (e.g. Dow 1981)^27^, the very tiny (<0.1°) receptive fields can be difficult to reliably characterize given ever-present ocular microsaccades. Furthermore, histological sectioning in anatomical studies may have incompletely preserved these lateral cortical territories if not specifically targeted^28^. As a consequence, the foveolar cortex has likely been under-studied or partially overlooked in prior electrophysiological and anatomical studies. These anatomical and electrophysiological constraints further underscore the need for a reliable noninvasive framework for localizing foveolar representations with high precision.

We have thus turned to study in anesthetized monkeys, in which ocular microsaccades can be removed using paralytics and fine resolution mapping can be achieved with ultrahigh field MRI. Here, we describe a successive spatial refinement strategy for mapping foveolar cortex by identifying the representations of the (0,0) intersection between the horizontal and vertical meridian representations in early visual cortex. This approach transforms meridian representations—traditionally used to delineate areal boundaries—into precise functional landmarks for central visual mapping. This method also reinforces our previous report of multi-locus representation of foveola and foveolar core^26^.

## Results

### Strategy for foveolar mapping in anesthetized macaques

The standard approach for mapping eye position in anesthetized animals is by standard back-reflection onto a tangent screen. This method, however, can have inherent spatial error (e.g. 1 degree or more) and cannot readily be done while the animal is in the MRI bore. We therefore developed a retinotopic mapping strategy of sequential refinement to precisely localize the foveolar representation in anesthetized macaques (**Fig. 1**). This strategy leverages the well-established organization of vertical and horizontal meridian representations in early visual cortex and applies a structured coarse-to-fine visual stimulation paradigm to progressively refine meridian coordinates. In early visual cortex, these meridians constitute robust functional landmarks that define areal boundaries and visual field transitions. The functional identification of the intersection of the horizontal (HM) and vertical (VM) visual meridians serves to locate the cortical representation of the foveola.

**Fig. 1.**
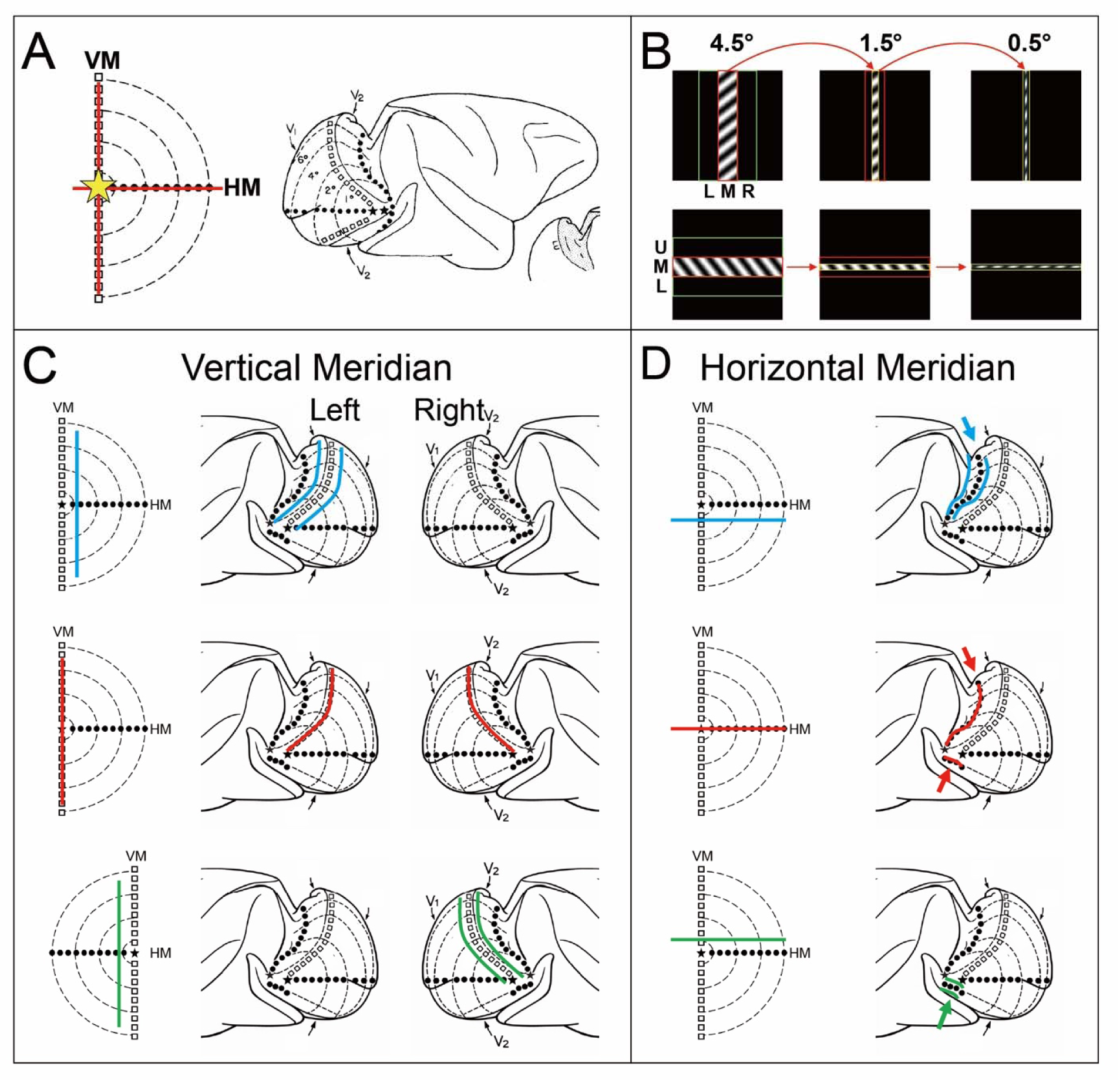
Strategy for functional localization of foveolar representation using sequential retinotopic refinement. (A) Visual field and cortical retinotopic organization. Schematic illustration of visual field coordinates and corresponding retinotopic organization in early visual cortex. The vertical meridian (VM; open squares) defines the boundary between V1 and V2, whereas the horizontal meridian (HM; filled circles) delineates the upper and lower visual field representations and forms the border between dorsal and ventral visual areas. Isoeccentricity contours illustrate increasing eccentricity away from the foveolar center. Modified from Gattass et al^29^. (B) Localizing by visual stimulus refinement. Retinotopic mapping was performed using 18 stimulus sets consisting of drifting grating bars of three spatial widths (4.5°, 1.5°, and 0.5° visual angle) presented at three adjacent spatial positions along each meridian axis. For VM and for HM, a coarse-to-fine refinement strategy was implemented, in which the centralmost bar location identified at each spatial scale was subdivided into three narrower bars (see text). (C) Mapping of vertical meridian. Vertical bar stimuli presented at 3 adjacent horizontal positions predict systematic shifts in cortical activation. Bars positioned within a single hemifield (blue, green) evoke activation within the contralateral hemisphere. Stimulus at the VM activates the V1/V2 boundary bilaterally (red). (D) Mapping of horizontal meridian. Horizontal bar stimuli presented at 3 adjacent vertical positions predict systematic shifts in cortical activation. Bars aligned with the HM activate both dorsal and ventral V2/V3 borders (red, arrowheads). Bars located below (blue) or above (green) the HM produce activation restricted to either dorsal or ventral V2/V3, respectively. Due to mirror-symmetric retinotopic organization across the V2/V3 border, non-meridian horizontal bars produce paired activations reflected across the V2/V3 border (blue, green). Note: Much of the ventral V2/V3 border is not visible in this view. This provides a functional signature for HM localization.

#### Coarse-to-fine VM mapping

The paradigm exploits the well-characterized layout of the meridians (**Fig. 1A**, white squares: VM, black dots: HM) and uses a successive refinement procedure to determine foveolar location (**Fig. 1B**). For each meridian, drifting grating bars (4.5°) were presented at three adjacent positions (total spanning 13.5°, green box). The position that induced the centralmost response (as determined by cortical location and largest cortical magnification) was selected (red box). This bar was then subdivided into 3 smaller bars (1.5°) and the centralmost one determined (yellow box). This centralmost location was divided into 3 even smaller bars (0.5°) and again the centralmost one determined (green box).

#### Vertical meridian mapping

For the VM, which is the location of the V1/V2 border, the predicted signature (**Fig. 1C**) is (a) a shift in cortical activation from one hemisphere (blue), to bilateral (red), to the other hemisphere (green) as the stimulus crosses the midline, and (b) a single line of activation at the V1/V2 border when centered and dual line activations (topographic reflections across the V1/V2 border) when off-center. This progression is documented in the data of Section 3.2 (**Fig. 2–3**).

**Fig 2.**
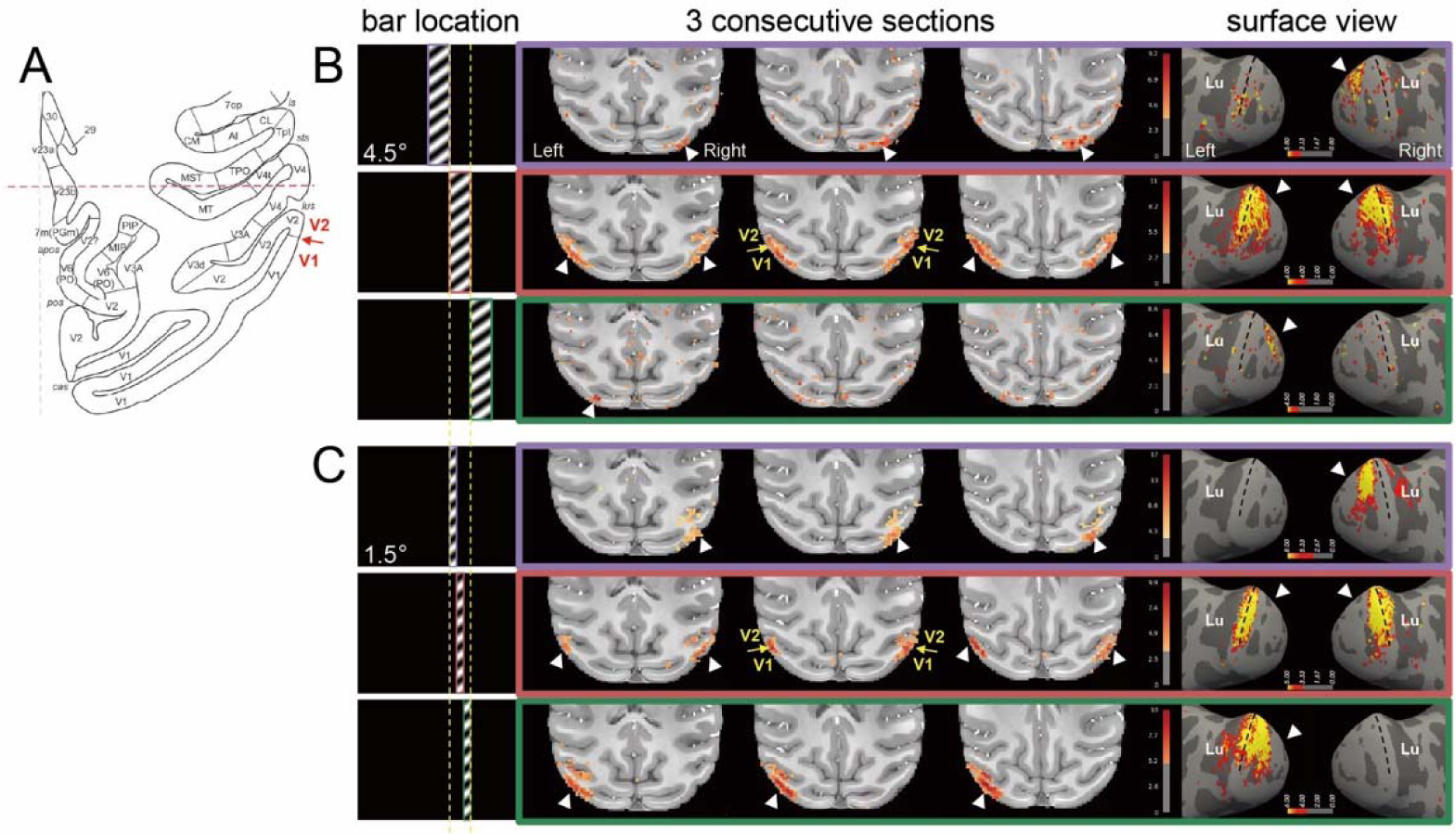
Determining visual vertical meridian with 4.5° and 1.5° wide bars. A. Predicted location of activation at V1/V2 border when stimulus overlies the VM (atlas slice adapted from Saleem & Logothetis]. B. Three 4.5° bars (with moving gratings embedded) were positioned at three adjacent locations (purple: upper row, red: middle row, green: lower row). When the bar covered the vertical meridian, robust and bilateral activations were seen at the V1/V2 border (middle row). Surface views shown in right column. C. At the selected middle bar position, three 1.5° bars were placed (yellow dashed lines). The bar overlying the VM activated the VM bilaterally (middle row), whereas off-center activations were shifted to the right hemisphere (purple) or to the left hemisphere (green). P<0.001. Monkey D.

**Fig 3.**
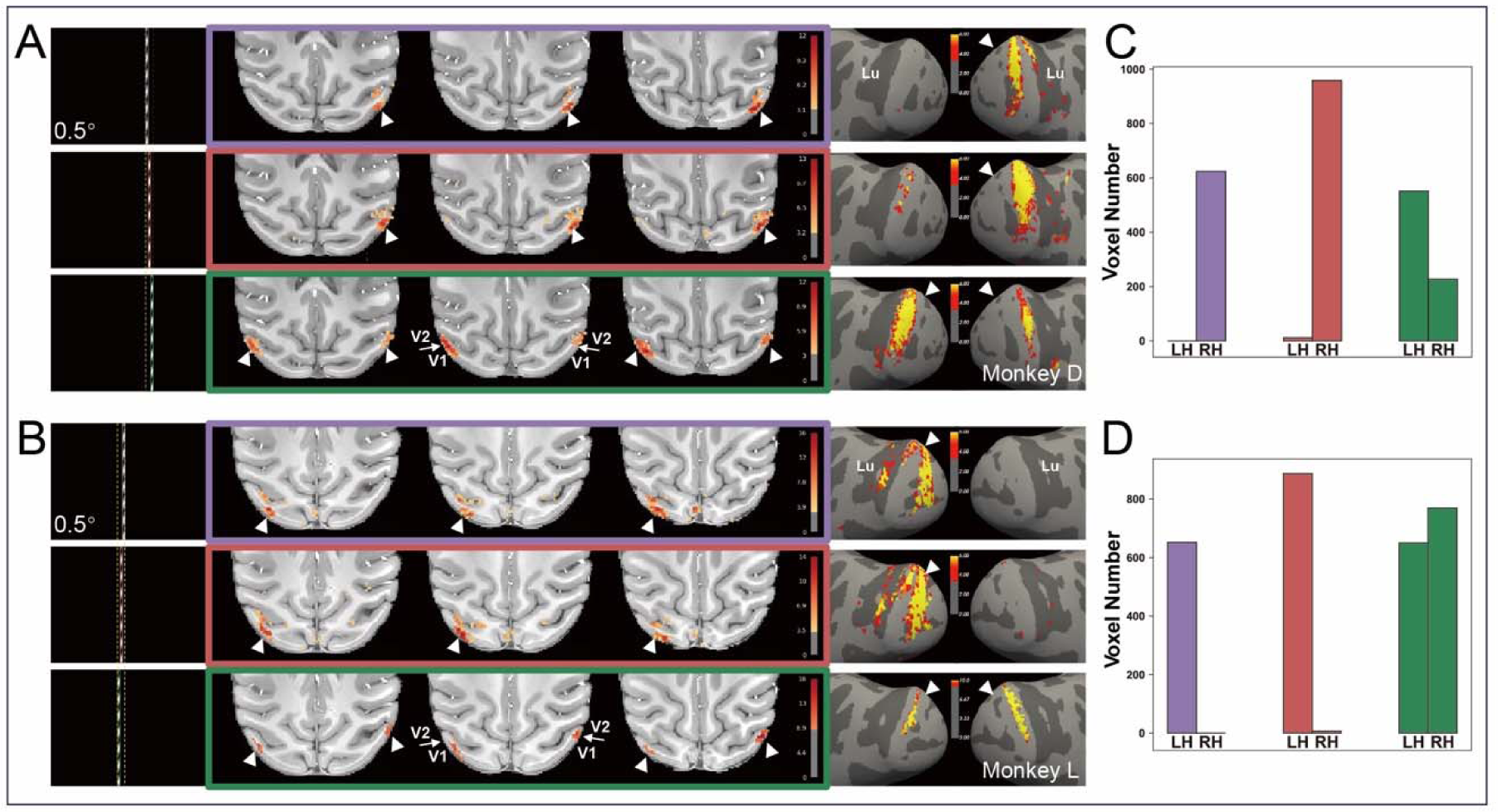
Vertical-meridian mapping using 0.5° wide vertical bars. (A, B) For each animal, 0.5° wide vertical bars were presented at adjacent horizontal positions (color-coded stimulus positions, shown at left, leads to activations shown in colored frames (purple, red, green) in middle panels. Slice views in middle panels show activation progressing from unilateral (A, B: white arrows in top two rows) or bilateral activations overlying the V1/V2 border indicating alignment of the stimulus position with the VM (A,B: white arrows in bottom row). Rightmost panels: Surface views show the corresponding transition from unilateral to bilateral activation. (C, D) Activated voxel counts in A, B, respectively, in the two hemispheres (LH, RH) for each bar position (purple, red, green). Bilateral activation (green) indicates position closest to foveolar X position. (A, C) Monkey D; (B, D) Monkey L. Statistical threshold: p < 0.001.

#### Horizontal meridian mapping

For the HM (**Fig. 1D**), horizontal bars were presented at three elevations. Because upper and lower fields are represented in both hemispheres, the HM is identified not by hemispheric asymmetry but by its dorsal–ventral signature: a bar on the meridian simultaneously activates both dorsal and ventral V2/V3 borders (red), whereas off-meridian bars below the HM drive only the dorsal cortex (blue) and above the HM drive only the ventral cortex (green, ventral HM extends to ventral surface and is only partially drawn). Similar to VM, there is a single line of activation at the V2/V3 border when centered (red arrowheads) and dual line activations (topographic reflections across the V2/V3 border) when off-center (blue and green arrowheads). Commonly, the HM falls in the fundus of the calcarine sulcus and serves as an anatomical landmark. This progression is documented in the data of Section 3.3 (**Fig. 4**).

**Fig 4.**
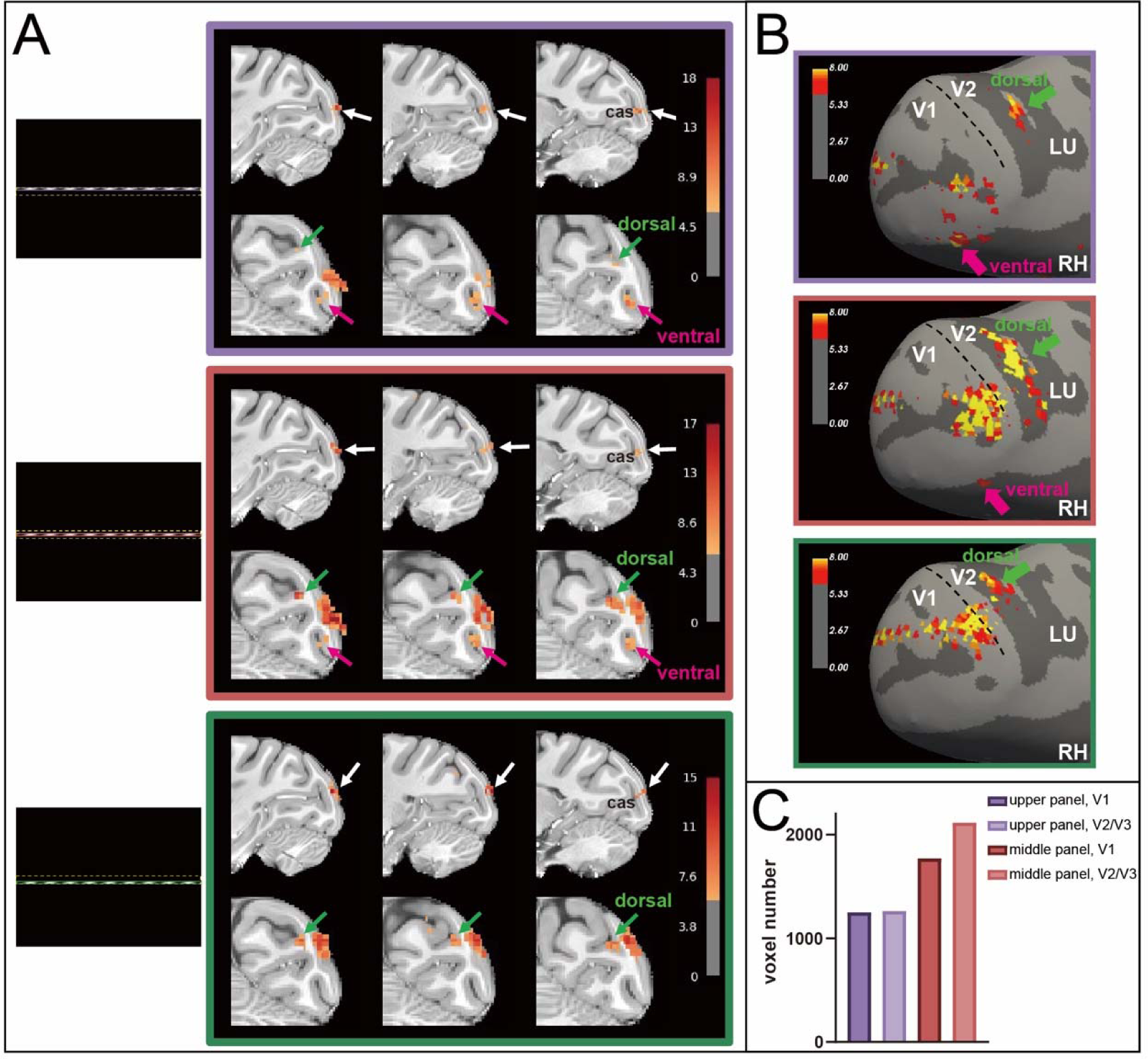
Horizontal meridian mapping with 0.5° wide bar. (A) Horizontal bars were presented at three adjacent vertical positions (upper, middle, and lower), producing a systematic shift of activation between ventral and dorsal portions of early visual cortex. 0.5° wide horizontal bars were presented at adjacent vertical positions (color-coded stimulus positions, shown at left) leads to activations shown in colored frames (purple, red, green). For each condition, sagittal views (upper row) and coronal views (lower row) are shown. White arrows indicate the location of the HM representation in V1, near the calcarine sulcus (labeled cas in the figure). Green arrows indicate dorsal V2/V3 activation and magenta arrows indicate ventral V2/V3 activation. The middle position (red) achieves the strongest dorsal and ventral activation, while the other two positions are heavily biased towards ventral (upper) or dorsal (lower). (B) Surface views (right hemisphere) for the same three conditions. Green arrows indicate dorsal V2/V3 activation and magenta arrows indicate ventral V2/V3 activation. The middle (red) condition activated both dorsal and ventral V2/V3 borders and produced the broadest, most spatially continuous response, and largest cortical magnification. LU, lunate sulcus; black dashed line, V1/V2 border. (C) Number of activated voxels in V1 and in V2/V3 for the upper (purple) and middle (red) conditions. Concomitant dorsal and ventral activation (red) indicates position closest to foveolar Y position. Monkey L. Statistical threshold: p < 0.001.

#### Foveolar mapping

The functional intersection of the independently refined VM and HM coordinates defines candidate foveolar coordinates (**Fig. 1A**, star). This is then further refined with localized 0.2° spot stimuli producing bilaterally symmetric activation across early visual areas (Section 3.4, **Fig. 5**). Together, this meridian-based hierarchical scheme localizes the foveolar representation under anesthesia and provides the foundation for the validation and functional analyses that follow.

**Fig 5.**
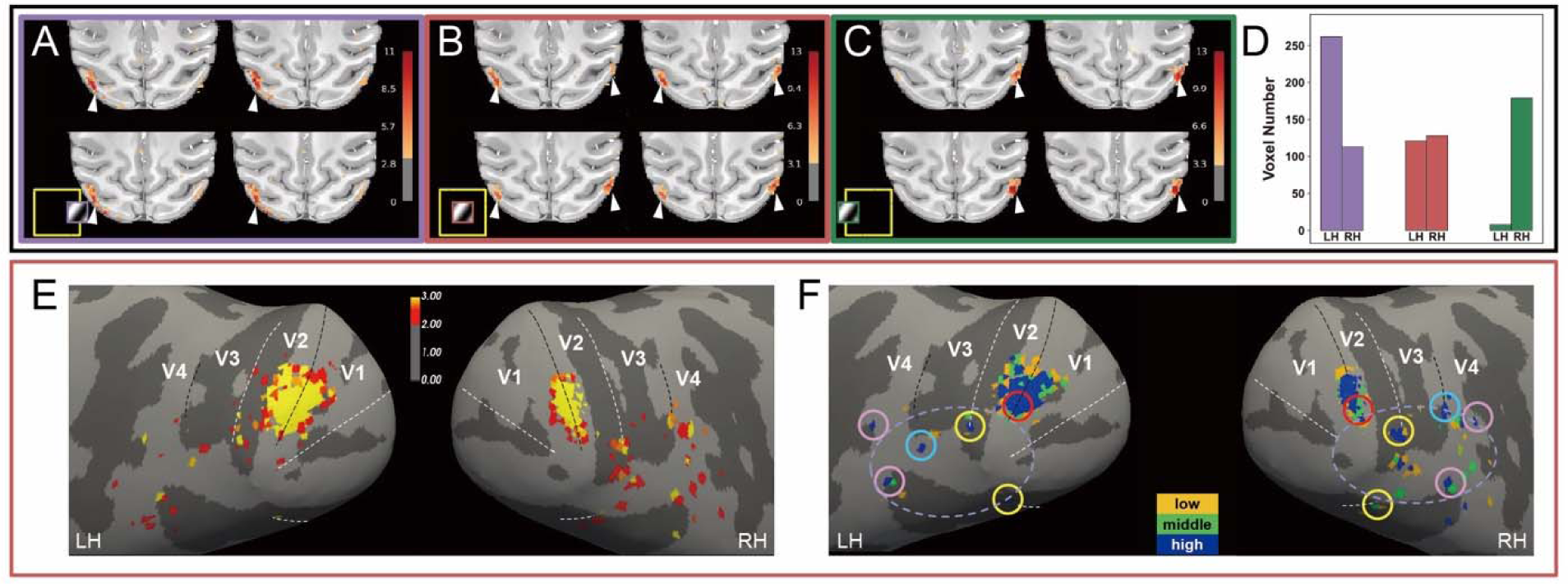
Refining foveolar localization using 0.2° spot stimuli. (**A–C**) Monkey L: activation evoked by three 0.2° spot stimuli presented within the 0.5° region defined by the intersection of the refined VM and HM coordinates — (**A**) spot displaced to the right of center (purple frame), (**B**) centered spot (red frame), and (**C**) spot displaced to the left of center (green frame); the three spots span 0.6° in total. The stimulus for each condition is shown in the yellow inset. For each condition, four horizontal slices through the center of activation are shown; arrowheads indicate activation evoked by the spot. A spot placed to the right of center activated mainly the left hemisphere (A) and a spot to the left mainly the right hemisphere (C), whereas the centered spot produced the most symmetric bilateral activation (B). (**D**) Number of activated voxels in the left (LH) and right (RH) hemispheres for each spot position; bar color corresponds to the frame color in A–C. (**E**) Surface projections of the activation evoked by the centered spot, shown for the left (LH) and right (RH) hemispheres. Dashed black lines mark the VM (V1/V2, V3/V4) and dashed white lines the HM (V1d/V1v, V2/V3). (**F**) Foveolar loci identified from threshold-dependent activation patterns (see Supplementary Fig. 2). Statistical thresholds are color-coded and normalized for each cortical area (low: yellow; middle: green; high: blue). Circles mark the foveolar loci (the lateralmost high-threshold activation for each area); note that, in left and right hemispheres, loci V1/V2v and V3/V4v are missing due to RF coil coverage resulting in lower SNR in ventral brain. The dashed purple oval marks the approximate perimeter of the foveolar core (note that the actual core trajectory follows gyral and sulcal contours). Statistical threshold for activation maps in A–C: p < 0.001.

### Vertical Meridian Mapping

Retinotopic activations exhibited systematic spatial shifts as a function of stimulus position (**Figs. 2-3**). Presentation of wide (4.5°) vertical grating stimuli at three adjacent horizontal locations produced distinct activation patterns in early visual cortex (see **Fig. 2A** for atlas-based slice reference). The stimulus position associated with the most extensive and bilaterally symmetric activation—consistent with representation of the visual midline—was identified as the candidate vertical meridian (VM) location (**Fig. 2B, middle row, red**). Shift of the bar leftwards (**top row, purple**) or rightwards (**bottom row, green**) produced relatively little and one hemisphere only activation.

Division of the middle row location into three narrower (1.5°) vertical gratings further refined VM localization. Of the three, the stimulus in the central position produced the strongest activation along the V1/V2 border and in both hemispheres (**Fig. 2C, middle row**). In contrast, stimuli off-center positions confined to one hemifield produced predominantly right hemisphere (**Fig. 2C, top row, purple**) or left hemisphere (**Fig. 2C, bottom row, green**) activation patterns and which were not centered on the V1/V2 border, consistent with retinotopic displacement away from the midline.

We further refined the mapping using 0.5° vertical gratings and show results from 2 monkeys. As shown in **Fig. 3** (Monkey D), as stimulus position shifted, cortical activation exhibited a systematic and continuous transition in hemispheric dominance. In the top row of **Fig. 3A (purple)**, activation is clearly confined to the right hemisphere. When the bar shifted 0.5° to the right (**middle row, red**), activation in the right V1/V2 border becomes more robust; in the left hemisphere, discontinuous activation patches likely corresponding to stripes in V2 are seen (surface view in right column). A further 0.5° shift to the right (**bottom row, green**), results in bilateral and continuous activations of the V1/V2 border (indicated by white arrows). Thus, while both the middle and bottom row exhibit bilateral activations, the bottom row exhibits clear continuous activation of the V1/V2 border in both hemispheres, and thus this stimulus location is considered most central of the three positions. An example from second monkey (Monkey L) is also shown in **Fig. 3B**. Again, the top two rows exhibit primarily single (right) hemisphere activation, which are not overlying the V1/V2 border. However, the bottom row (green) reveals bilateral activation which is over the V1/V2 border, identifying the centralmost location

Quantitative analysis further supports these observations. We quantified the number of voxels in the left (LH) and right (RH) hemispheres for each of the grating stimuli positions. For each of Monkey D and Monkey L, only one position (the rightmost of the 3 for each monkey) produced strong bilateral activation. The balance in voxel count between hemispheres, a quantifiable localization index, provides an objective and reproducible metric for vertical meridian identification.

### Horizontal Meridian Mapping

As will be shown below, the HM can also be defined by a characteristic pattern of simultaneous activation in both dorsal and ventral divisions of extrastriate cortex. Anatomically, the HM location is known to be in or near the fundus of the calcarine sulcus^22,29^, dividing the primary visual cortical operculum into dorsal and ventral representations of the inferior and superior visual fields, respectively. In addition, HM lies on the V2/V3 border, with both a dorsal and ventral representation; a shift into the superior visual field will activate only cortex near the ventral V2/V3 border and into the inferior field will activate only near the dorsal V2/V3 border. A stimulus that falls exactly on the HM will have partial activation in both dorsal and ventral V2/V3 cortex. A key defining aspect is that a line on the HM will activate foveal cortex with high cortical magnification.

A hierarchical mapping strategy analogous to that for VM was applied to localize the horizontal meridian (HM), consisting of progressive refinement from 4.5° to 1.5° (not shown) to 0.5° gratings. **Fig. 4A** & **4B** show the result of activation with the identified 0.5° gratings.

Functionally, horizontal gratings positioned away from the HM preferentially activated either dorsal or ventral V2/V3, consistent with stimulation of either the lower or upper visual-field (**Fig. 4A**, arrows; **4B**). As the stimulus approached the HM, activation began to involve both dorsal and ventral V2 & V3 cortex (white arrows in **Fig. 4B** upper and middle panels), indicating a well centered horizontal stimulus recruiting representations on both sides of HM. This near-HM pattern was visible in **Fig. 4B** upper panel, where activation exhibits some contact with the HM (white arrows). In contrast, the middle row produced the broadest and most spatially continuous V2/V3 border activation, indicating that this stimulus position straddled the horizontal meridian most fully (**Fig. 4B**, middle row; white arrow). Quantification of the activated voxels in V1 and V2/V3 supported this interpretation: of the tested vertical positions, the HM-aligned condition elicited the largest activation (red bars), thereby providing a quantitative way to identify the horizontal-meridian location (**Fig. 4C**). Note that the location of the HM in V1 is coincident with the calcarine sulcus (white arrow). Similar results were found in a second monkey (**Fig. 4 Monkey L, Supplementary Fig.1 Monkey D**). This dorsal–ventral transition is consistent with the established retinotopic organization of early visual cortex and also consistent with the fundus location of the calcarine sulcus location, providing both anatomical and functional landmarks for horizontal meridian localization.

### Bilateral Foveolar Representation

Following independent localization of the VM and HM, their intersection was used to further refine foveolar coordinates. As shown in **Fig. 5A-C (Monkey L, for Monkey D see Supplementary Fig. 3)**, centered on the 0.5° intersection region (yellow frame), three small spot stimuli (0.2° diameter) were presented: one 0.2° spot centered on the x, y coordinate from above fine 0.5° width grating meridian mapping (red outline, **Fig. 5B**), and two 0.2° adjacent spots to the right (purple outline, **Fig. 5A**) and to the left (green outline, **Fig. 5C**), with a total horizontal extent spanning 0.6°. **Fig. 5A–B** (Monkey L) reveals that systematic shifts in spot position leads to predictable shifts in hemispheric activation patterns (four central horizontal slices through the centers of activation are shown for each spot position). When the stimulus was centered at the candidate foveolar location, strong bilateral BOLD responses were observed (**Fig. 5B**). When the spot was positioned to the right (activating the right hemi visual field), activation shifted primarily to the left hemisphere (**Fig. 5A**) and when the spot was positioned to the left, activation shifted primarily to the right hemisphere (**Fig. 5C**). As the spot position in **Fig. 5B** produces the most balanced bilateral activation (quantified in **Fig. 5D**, it is thus the centralmost foveolar position. This refinement further demonstrates localization of the foveolar representation to within 0.2° precision in anesthetized monkeys.

Projection of activation maps onto reconstructed cortical surfaces revealed discrete foveolar loci distributed along early visual-area borders (**Fig. 5E**). Note correspondence with VM (V1/V2, V3/V4) and HM (V1d/V1v, V2/V3) meridia (dashed black and white lines, respectively).

Threshold-dependent analysis further identified multiple candidate foveolar loci, typically centered on lateral ends of visual-area borders (**Fig. 5F circles;** see method **Supplementary Fig. 2**). To identify the most foveolar activation, we selected, for each area, the lateralmost high threshold activation. We recognized that different areas respond with different robustness to simple flashing spot stimuli; so we employed normalized threshhold ranges for each cortical area (**Fig. 5F**, Blue: high threshold; Green: middle threshold; Yellow: low threshold)^26^. This revealed the very laterally located blue voxels (colored circles in **Fig. 5F**); the most lateral foci with high threshold that remained spatially stable across thresholds were taken as candidate foveolar loci (**Supplementary Fig. 2**). Note that, due to the placement of the surface coil over dorsal visual cortex, activation in some ventral visual areas was not consistently detected (see **Supplementary Fig. 4** coil coverage signal map). Despite this (as shown in our previous study in awake fixating monkeys), the locations are consistent with ventral and dorsal V1/V2 (red), V2/V3 (yellow), V3/V4 (blue), and V4/TEO (pink) foveolar loci. These loci define the perimeter of the newly described ‘foveolar core’(dashed purple oval)^26^. This distribution is consistent with prior observations in awake macaques and supports the presence of a specialized cortical territory associated with central visual processing.

### Location of foveolar loci across animals

In surface view, foveolar loci were identified using the same threshold-dependent procedures described above^26^ (**Fig. 5F; Supplementary Fig. 2**) on the inflated cortical surface of both hemispheres and numbered 1–8 according to their positions along early visual-area borders of Monkey L (**Fig. 6A**). The dashed lines mark the V1/V2, V2/V3, and V3/V4 boundaries. Loci 1 and 2 lie closest to the common foveal endpoint of V1 and V2; the remaining numbered loci form progressively more lateral and inferior pairs along successive early visual-area borders toward V4/TEO. The same color-number pairs occupy corresponding positions in the two hemispheres, producing an approximately mirror-symmetric sequence.

**Fig 6.**
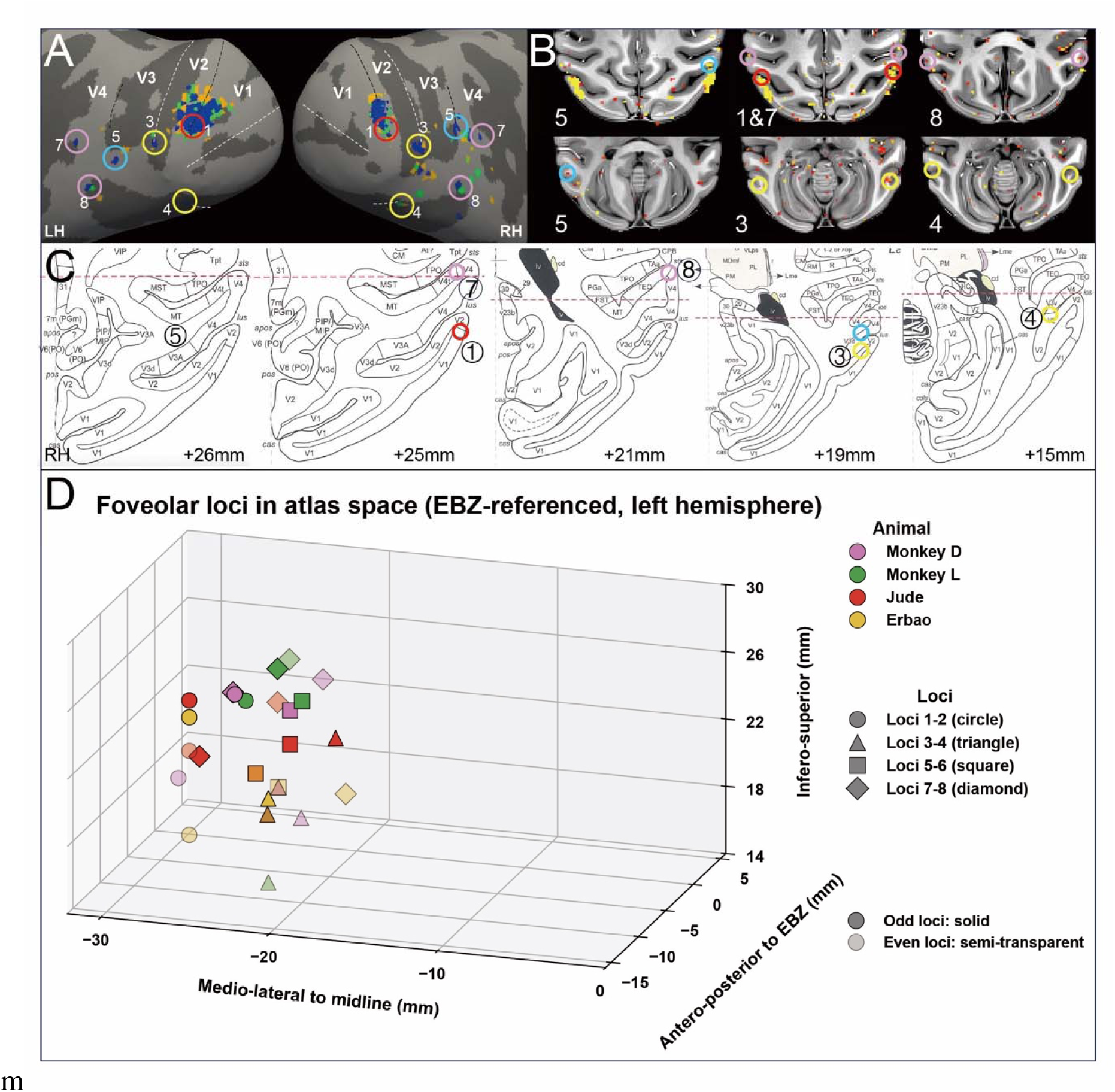
Spatial localization of foveolar cortical loci across surface, anatomical, atlas, and coordinate spaces. (A) Foveolar loci shown on the inflated cortical surface of the left (LH) and right (RH) hemispheres of Monkey L, numbered 1–8 according to their positions along the borders of early visual areas (V1/V2, V2/V3, V3/V4, and V4/TEO). Dashed lines mark the areal borders. (B) Representative foveolar loci shown on axial (horizontal) anatomical MRI sections; labels and coloured circles denote individual loci (upper row: 5, 1&8, 7; lower row: 5, 3, 4). Monkey L. (C) Individual foveolar loci localized on the corresponding plates of a standardized macaque atlas (Saleem & Logothetis), shown at successive infero-superior (dorsal) levels relative to the ear-bar zero (EBZ)^34^ reference plane (+26, +25, +21, +19, and +15 mm; dashed red line indicates the reference level). Circled numbers indicate individual loci (right hemisphere). (D) Distribution of foveolar loci from four animals in EBZ-referenced atlas coordinate space, plotted for the left hemisphere in three dimensions (medio-lateral distance to midline × antero-posterior distance to EBZ × infero-superior distance to EBZ). Symbol colour denotes animal — Monkey D and Monkey L (anesthetized, this study) and Monkey J and Monkey E (awake, previous study) — and symbol shape denotes locus group (loci 1–2, circle; 3–4, triangle; 5–6, square; 7–8, diamond); odd-numbered loci are plotted solid and even-numbered loci semi-transparent. Across animals, foveolar loci were confined to the most lateral occipital cortex, ∼18 to 30 mm from the midline, spanning approximately −10 to +4 mm along the antero-posterior axis (caudal–rostral relative to EBZ) and +15 to +26 mm dorsal (infero-superior) to EBZ.

In axial anatomical sections (**Fig. 6B**), foveolar responses appeared as multiple discrete activation clusters distributed along the borders of the early visual areas (V1/V2, V2/V3, V3/V4, and V4/TEO). This multi-focal pattern reproduces, in anesthetized animals, the organization we previously described in awake, fixating macaques, in which the foveola is represented by discrete loci rather than a single confluent focus^26^. The present clustering confirmed that this distributed, multi-locus organization is also present under anesthesia.

A key question is whether these foveolar loci occupy consistent anatomical positions across animals and hemispheres, or whether their locations vary substantially between individuals. We found that, across eight hemispheres (four from previous paper in awake monkeys^26^,four from this study in anesthetized monkeys), the loci occupied comparable positions along the V1/V2, V2/V3, V3/V4, and V4/TEO borders, forming an approximately mirror-symmetric arrangement about the midline that was reproducible across animals. To relate these functional activations to anatomical coordinate space, each foveolar locus was localized on the corresponding slice of a standardized macaque atlas^30^ (**Fig. 6C**). This atlas was chosen because it provides both histologically validated areal boundaries and MRI images in a common stereotaxic frame referenced to the ear-bar zero (EBZ), allowing loci from different animals to be compared in a shared coordinate space. Reading the atlas planes from dorsal to ventral, locus 5 appears near +26 mm, loci 1 and 7 near +25 mm, locus 3 near +21 mm, locus 3 near +19 mm, and locus 4 near +15 mm relative to EBZ. Reproducibility test across four animals were shown in **Fig. 6D** by pooling the EBZ-referenced coordinates from two awake animals (Monkey E and Monkey J)^26^ and two anesthetized animals (Monkey L and Monkey D). Symbol color identifies the animal, and symbol shape identifies loci 1-8. The points form left hemisphere lateral-occipital loci approximately 18-30 mm from the midline. Most loci lie from about 10 mm caudal to 4 mm rostral to EBZ; within each hemisphere, points with the same shape recur in similar relative positions despite differences among animals. In the posterior projection, most loci occupy a band approximately 15-26 mm dorsal to EBZ and again form lateral pairs. Thus, the principal result is not that there is approximate bilateral symmetry. The overlap of awake and anesthetized cases further indicates that this distributed organization is not created by anesthesia. Together, the panel-by-panel evidence supports a multi-loci ‘foveolar core’ whose topology is reproducible even though its exact coordinates vary among hemispheres and individuals.

### 3.6 Location of the foveolar core

As the foveolar loci are localized with the method described above, we now define the ‘foveolar core’. To make this territory explicit, the loci were connected in surface view (yellow outline, **Fig. 7A**). In both hemispheres the connected loci formed a closed, region spanning the V1/V2, V2/V3, V3/V4, and V4/TEO borders at the most lateral extent of the visual cortex.

**Fig 7.**
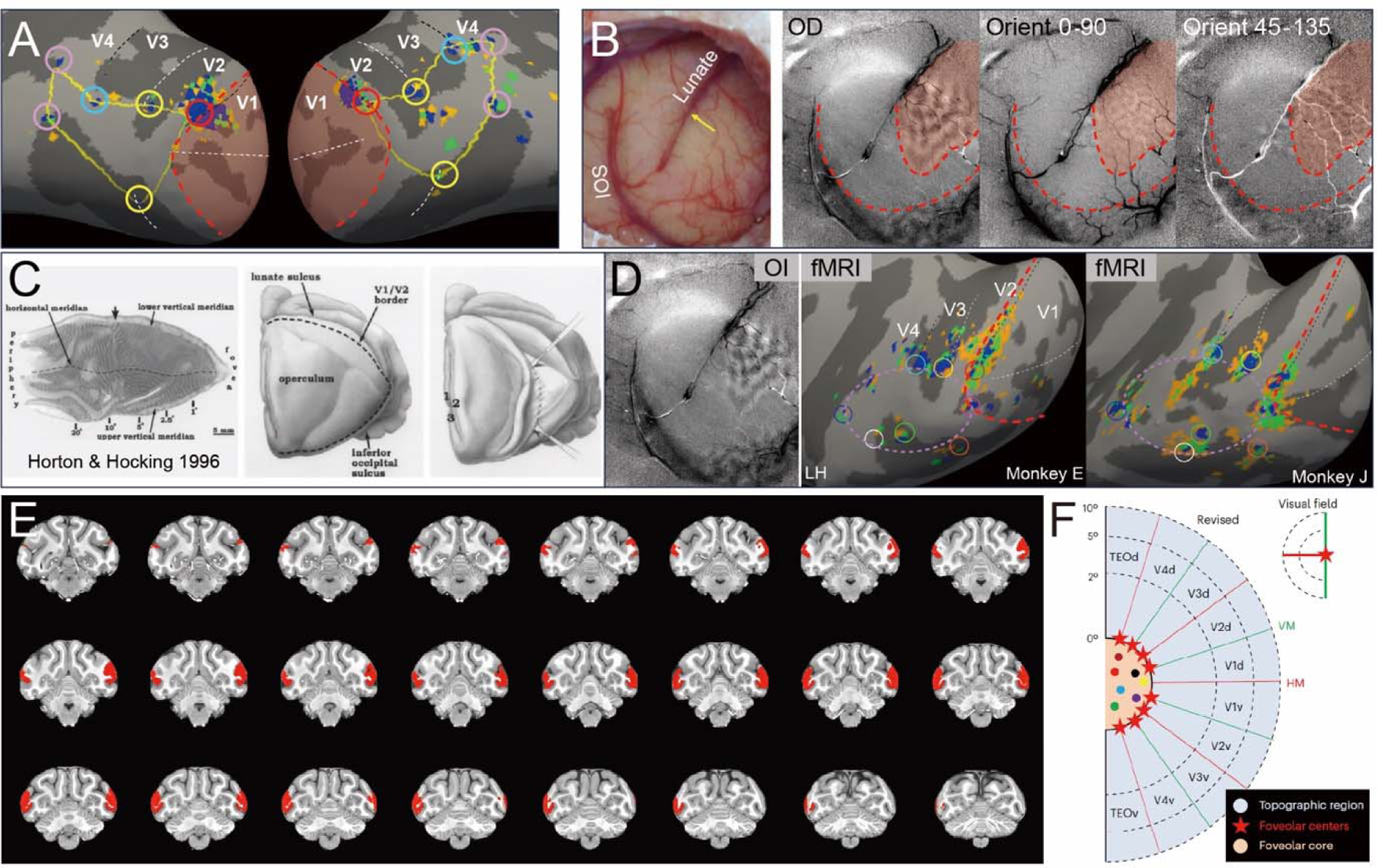
The foveolar core is a distinct area. (**A**) Foveolar loci (colored circles) shown on the inflated left and right hemispheres and connected in sequence (yellow lines) to delimit the ‘foveolar core’. Black and white dashed lines mark the V1/V2, V2/V3 and V3/V4 borders; the red dashed contour and red shading indicate the extent of V1. (**B**) Intrinsic-signal optical imaging of the lateral operculum. Left: blood-vessel map of the imaged field showing the inferior occipital sulcus (IOS) and the end (yellow arrow) of the lunate sulcus (LUN). Right: ocular-dominance (OD) and orientation difference maps (0–90°; 45–135°) obtained from the same field. These functional OD and orientation maps are clear in V1, but ends abruptly at the border with the core (red dashed line). The core is bounded by dorsal V1 and ventral visual areas (delimited by red dashed lines), between which the core lies (red shading). (**C**) For comparison, flat-mounted and stained primary visual cortex from Horton & Hocking (1996), showing the horizontal, upper and lower vertical meridian representations and the fovea-to-periphery axis (left), and the extent of the excised and flattened tissue relative to the lunate sulcus, the V1/V2 border, the operculum and the inferior occipital sulcus (middle, right). (**D**) Alignment of the excision boundary of Horton & Hocking (black dashed line) with the V1 border (red dashed lines) in our own optical imaging (OI, left) and fMRI surface maps (middle: Monkey E, left hemisphere; right: Monkey J), showing that the excised tissue did not include the core region. Colored circles indicate foveolar loci as in A. (**E**) The core outlined in A seen in successive coronal sections through the occipital lobe (red). Colored circles indicate individual foveolar loci, colour-matched to A. The projected core forms a continuous territory along the lateral surface across successive coronal planes. (**F**) Schematic of summary for the organization of foveolar loci and foveolar core from 4 monkeys: eight foveolar loci (eight red stars), one at each of the dorsal and ventral representations of V1/V2, V2/V3, V3/V4 and V4/TEO; the foveolar core (orange) is an area within the ring of stars and outside the area of visuotopic representation.

Consistent with the ordered sequence, the outline was obtained in each hemisphere by following the same locus order. We refer to this bounded territory as the ‘foveolar core’, as in our previous study in awake, fixating macaques^26^.

Previously, we had shown that the core contains functional domains sensitive to high spatial frequency stimuli, color stimuli, and motion stimuli but was not responsive to small stimuli and not topographic by phase encoding mapping methods, raising the possibility that the core is a higher order area^26^. Here, we further show using intrinsic-signal optical imaging that the core does not respond to standard visual gratings (**Fig. 7B**). The imaged field is located over the lateral operculum near the inferior occipital sulcus (IOS) and the lunate sulcus (blood-vessel map, left). In the adjacent retinotopically organized cortex, the difference maps resolved the expected columnar architecture — ocular-dominance bands (OD) and orientation domains (0–90°; 45– 135°). Interestingly, these functional maps did not extend into a region near the IOS and the end of the Lunate sulcus and instead ended abruptly (dashed red curved line), appearing uniformly smooth, with no comparable ocular-dominance or orientation domain structure. This lack of response helps to further define the location of the core. It is consistent with potential higher order nature of the foveolar core and provides further evidence for its distinctions from traditional areas in early visual cortex.

To provide additional understanding of where this core area is, we compared this to well-known studies by Horton and Hocking in which primary visual cortex was excised in its entirety, flattened, and stained to reveal ocular dominance maps (**Fig. 7C**). Aligning the excision boundary (**Fig. 7C**, black dashed line) with the V1 boundaries (red dashed lines) in our brain maps as seen in OI (**Fig. 7D**, left) and fMRI (**Fig. 7D**, middle, right) reveals that the excised tissue did not include the core region.

Finally, to place this surface territory in the same volumetric frame used above, the core outlined in **Fig. 7A** is shown on coronal sections through the occipital lobe (red, **Fig. 7E**); circled loci are the individual foveolar loci, color-matched to **Fig. 7A**. The projected core is thus seen as a continuous territory along the lateral surface across successive coronal planes, with individual loci appearing at discrete antero-posterior levels around its perimeter. This is the volumetric counterpart of the atlas-space distribution in **Fig. 6**: the same region bounded by foveolar loci are seen here as sequential coronal volume slices.

Together, these observations extend the multi-locus description of **Fig. 6** in two ways. First, the foveolar loci are not isolated activation foci but the boundary of a definable cortical territory that can be identified in surface, optical, and volumetric views. Second, the core territory is not simply a continuation of the surrounding early visual cortex: it differs from the columnar architecture that characterizes adjacent topographic areas and indicates that the ‘foveolar core’ is a functionally distinct area (**Fig. 7F**).

## Discussion

### Defining foveolar gaze in anesthetized monkey with precision

As part of an effort to investigate foveolar vision, we recently reported that the ‘foveolar confluence’ of the macaque monkey comprises multiple foveolar representations (8 per hemisphere) encircling a non-visuotopic region we term the ‘foveolar core’. This study was conducted in awake monkeys trained to perform precise fixation in a 7T MRI. However, to study foveolar cortex in anesthetized animals in the MRI, new methods are needed to locate the anesthetized animal’s center of gaze on a monitor with high precision. Here, we established systematic methods to precisely localize (to within 0.2°) the location of foveolar gaze, thereby introducing a critical step that enables further study of foveolar cortical function. Specifically, we developed a meridian-based (HM and VM) retinotopic mapping strategy that systematically presented coarse-to-fine vertical and horizontal bars and spot stimuli. Identification of the HM and VM intersection thus provided the foveolar locations with sub-degree spatial precision. Moreover, vertical meridian was identified by bilaterally symmetric activation across hemispheres, and horizontal meridian localization by simultaneous dorsal and ventral activations (**Fig. 2-4**). Together, these signatures provided convergent functional and anatomical criteria for meridian identification.

Previous studies have established fMRI as a powerful tool for mapping visual cortical organization in nonhuman primates, both in awake and anesthetized conditions. Classical retinotopic paradigms, including rotating wedges, expanding rings, and sweeping bars, have been highly effective in delineating large-scale visual field organization and defining boundaries between early visual areas such as V1, V2, and V3. However, these approaches are primarily optimized for mapping broad retinotopic gradients and are not specifically designed to resolve the centralmost visual field with sub-degree precision. In awake, behaving monkeys, precise foveal mapping is facilitated by fixation control and eye tracking^15,31,24,32^. In contrast, anesthetized preparations provide highly stable imaging conditions suitable for ultra-high-resolution fMRI, and integration with invasive recording and stimulation methodologies. Even though paralytic is used, the possibility of slow eye drift cannot be ignored; methods of conduct precision monitoring of fine eye movements (e.g. using a Purkinje eye tracker^33^) in the MRI environment remain to be developed. Despite these factors, our results demonstrate that such eye drift under anesthesia is minimal and that accurate foveolar localization can nevertheless be achieved by leveraging robust stimulus-driven retinotopic signatures (**Fig. 1**) in ultrahigh field MRI.

### The foveolar core: defining a new visual area

Beyond the mapping strategy itself, this study provides the first stereotaxic definition of the foveolar core. These methods also further confirmed, in anesthetized macaque monkeys, the presence of multiple foveolar representations and ‘foveolar core’ in each hemisphere. Pooling loci from eight hemispheres of four animals into a common ear-bar-zero frame^34^ (**Fig. 6**) places them within a restricted territory of the most lateral occipital cortex, ∼18–30 mm from the midline, −10 to +4 mm antero-posterior and +15 to +26 mm dorsal to EBZ. The data also illustrate some degree of variability across individuals. To further help the reader locate this region, we also provide optical imaging views of this cortical area by illustrating its lack of response to classical gratings. This provides clearly visible borders of this region, nestled near the IOS and the end of the lunate sulcus (**Fig. 7B**). Practically, this provides both cortical surface views, coronal/saggital/horizontal views of sulcal locations, as well as anatomical coordinates to enable placement of electrodes, tracer injections, or optical windows of this new region.

Why was such a region not recognized earlier? While it is possible that some studies may have recorded from this location^35^, its extreme lateral location and vertical orientation of the cortex inferior occipital sulcus, made it difficult to access with standard vertical chambers and electrode penetrations. Moreover, the very small receptive fields (<0.1°) of foveolar neurons are confounded by ever-present microsaccades, making electrophysiological characterization difficult. In addition, as shown in (**Fig. 7C, D**), the classical flatmount preparations that famously illustrated the architecture of excised V1 tissue excludes this foveolar core territory^28^. Thus, for multiple reasons this region remained understudied and unrecognized.

This carries consequences for how the early visual map is understood. In the classical account, V1–V4 form a continuous topographic mosaic converging on a single foveal confluence^21^. Our data indicate that the map is instead interrupted at its centre: the confluence is not a point of convergence but a ring of discrete loci surrounding a territory that lacks the ocular-dominance and orientation columns of the areas around it (**Fig. 7F**). The foveolar representation is therefore not simply the high-magnification end of the early visual areas, but is accompanied by a separate, apparently higher order cortical area. Its function remains to be elucidated.

In summary, the present study establishes a robust and reproducible strategy for foveolar localization in anesthetized macaques. This capability opens multiple new avenues for study. For example, INS-fMRI (a method using focal stimulation in ultrahigh field MRI to map brainwide mesoscale networks^36^) can be used to understand the connectivity of foveolar core with brainwide motor, parietal, and prefrontal areas. As well, structural and functional laminar contributions to feedforward and feedback connectivity can be determined^38^. Application of exciting methods for mapping cortical microvascular *in vivo* may further our understanding of hemodynamics and energetics of foveolar cortical function in health and disease^37^. Importantly, this approach provides a solid basis to evaluate functional responses in monkeys performing behavioral tasks in the MRI.

## Methods

### Animal Preparation

All experimental procedures conformed to the National Institutes of Health Guide for the Care and Use of Laboratory Animals and were approved by the Institutional Animal Care and Use Committee of Zhejiang University.

Two adult macaque monkeys (Monkey D: 4 years old, 5–6 kg, male; Monkey L: 6 years old, 6–7 kg, female) were used in this study. Animals were initially sedated with ketamine hydrochloride (10 mg/kg, intramuscular) and atropine sulfate (0.03 mg/kg, intramuscular).

Animals were subsequently intubated, mechanically ventilated, anesthetized with 1%–2% isoflurane in oxygen, and head secured in a custom-built MR-compatible stereotaxic frame. Physiological parameters, including core body temperature (maintained at 37.5–38.5°C), heart rate, respiratory rate, arterial oxygen saturation (SpO), and end-tidal CO, were continuously monitored throughout the experiment.

During functional imaging, anesthesia was maintained using continuous intravenous infusion of sufentanil (induction dose 3 μg/kg; maintenance 2–6 μg/kg*h) supplemented with low-dose isoflurane (0.2%–0.5%). To eliminate eye drift which can still occur under anesthesia, neuromuscular blockade was achieved using vecuronium bromide (induction dose 0.3 mg/kg; maintenance infusion 0.1 mg/kg*h)^36,38,39^. As the eyes are not aligned under anesthesia, visual stimulation was delivered to one eye while the contralateral eye was occluded using opaque blackout material; this ensured unilateral visual input.

### Eye Position Estimation

At the beginning of each session, the refractive state of each eye was determined and corrected so that the stimulus monitor lay in the plane of best focus. Following pupil dilation and cycloplegia (topical 1% atropine), refraction was assessed by streak retinoscopy at the experimental viewing distance. Rigid gas-permeable contact lenses of the power required to bring the retina into focus on the screen were then fitted to each eye, both to correct refractive error and to protect the cornea throughout the experiment.

Because the experiment was conducted inside the MRI scanner, ferromagnetic instruments such as the fundus ophthalmoscope could not be brought into the magnet room, precluding direct back-projection of retinal landmarks onto the stimulus screen. The initial estimate of the animal’s line of sight was therefore obtained by visual inspection of the eye and its gaze direction by the experimenter, which provides only an approximate indication of the foveal projection on the monitor. To ensure that this coarse estimate did not cause the foveal representation to be missed, the first stage of mapping deliberately used wide-field stimulation. Three 4.5° bars were presented at adjacent positions spanning a total of 13.5° across the visual field, so that the central line of sight was reliably encompassed within the stimulated region regardless of the residual uncertainty in the initial gaze estimate. This wide coverage guaranteed that the foveal representation fell within the activated cortical territory, which was then progressively narrowed through the subsequent coarse-to-fine refinement.

### Visual Stimuli

To localize the vertical meridian (VM) and the horizontal meridian (HM), retinotopic mapping was performed by functional mapping in response to a series of vertical and horizontal gratings, respectively, presented in a coarse-to-fine refinement strategy. For each of VM and HM, 3 sets of 3 vertical and horizontal grating conditions (3 widths × 3 positions) were used, respectively, resulting in nine stimulus conditions per meridian and a total of 18 stimulus conditions across both meridians. Each meridian stimulus consisted of drifting sinusoidal gratings (spatial frequency: 3 cycles/degree; temporal frequency: 2 Hz; orientations: 45° and 135°, each presented for 10 s). Visual stimuli were generated using VPixx software (VPixx Technologies) and projected onto a rear-projection screen positioned 57 cm from the animal’s eyes using a PROPixx projector (1920 × 1080 resolution; 60 Hz RGB refresh; 1440 Hz grayscale mode).

For VM localization, grating stimuli were presented over the estimated foveal location on the monitor at three adjacent positions spanning the horizontal visual axis (left, center, and right).

Initial localization of central visual responses was performed using wide (4.5°) grating to identify a candidate VM position. The selected vertical grating position was divided into 3 narrower gratings (1.5°) and the best VM position selected. This best position was again divided into 3 even narrower gratings (0.5°), after which the best (and final) VM position was determined.

Similarly, for HM localization, grating stimuli were presented at three positions spanning the vertical visual axis (upper, middle, and lower), and a coarse-to-fine (4.5°, to 1.5°, to 0.5°) localization was performed.

In this manner, VM and HM coordinates were independently refined at the highest (0.5°) spatial resolution, and their intersection used to define the foveolar X,Y coordinates. Final determination of the V1/V2, V2/V3, V3/V4, and V4/TEO foveolar activation centers was performed using a 0.2° diameter grating stimulus presented at the determined X,Y coordinates. Bilateral symmetry of cortical activation was used as a validation criterion.

### MRI Acquisition and Preprocessing

All MRI data were acquired using a 7 Tesla Siemens Magnetom scanner. High-resolution structural images were obtained under isoflurane anesthesia using a three-dimensional MPRAGE sequence with the following parameters: repetition time (TR) = 2590 ms; echo time (TE) = 2.73 ms; inversion time (TI) = 1050 ms; matrix size = 192 × 192; field of view (FOV) = 96 × 96 mm²; slice thickness = 0.5 mm; flip angle = 7°; bandwidth = 250 Hz/pixel; number of averages = 3; total acquisition time = 24 min 52 s.

Functional images were collected using a 16-channel surface coil optimized for visual cortex imaging^40^. Blood oxygenation level-dependent (BOLD) contrast images were acquired using single-shot T2*-weighted gradient-echo echo-planar imaging (GRE-EPI) with the following parameters: TR/TE = 2000/35 ms; matrix size = 182 × 140; FOV = 110 × 85 mm²; voxel size = 0.8 mm isotropic; flip angle = 70°; bandwidth = 784 Hz/pixel; echo spacing = 1.44 ms; GRAPPA acceleration factor = 3.

Stimuli were presented using a block design consisting of alternating blank and stimulus periods (20 s blank followed by 20 s stimulation composed of two 10 s grating orientations). Each run contained 10 stimulus blocks and lasted approximately 7 minutes.

### Online and Offline fMRI Analysis

Real-time activation maps were generated using Siemens integrated BOLD analysis software. Activation maps were overlaid onto anatomical images and visualized using Neuro 3D software. Activation maps were thresholded at q < 0.05 after FDR correction; for visualization, maps are shown using an equivalent voxel-level threshold of p < 0.001.

Offline preprocessing and statistical analyses were performed using AFNI and FreeSurfer (version 6.0). DICOM files were converted to NIfTI format using AFNI’s Dimon utility.

Functional images were reoriented from supine to sphinx position and underwent slice timing correction, susceptibility distortion correction using reverse phase-encoded images, and rigid-body motion correction. Spatial smoothing was intentionally omitted to preserve high spatial resolution^41^.

Functional volumes were projected onto cortical surface reconstructions using landmark-based registration^42,43^. General linear model (GLM) analyses were conducted using AFNI’s 3dDeconvolve^44^, incorporating stimulus timing convolved with a canonical gamma hemodynamic response function (HRF) and six motion parameters as nuisance regressors^45^. Low-frequency signal drift was removed using fifth-order polynomial detrending^46^.

### Ethics statement

All experimental procedures were conducted in accordance with the National Institutes of Health Guide for the Care and Use of Laboratory Animals and were approved by the Institutional Animal Care and Use Committee of Zhejiang University. All efforts were made to minimize animal discomfort and to reduce the number of animals used.

## Data availability

The full 7T fMRI and intrinsic-signal optical imaging datasets generated in this study are available from the corresponding authors on reasonable request; restrictions apply to the raw imaging data owing to their size. The macaque atlas used for stereotaxic referencing is available from its original publication.

## Code availability

This study used the publicly available packages AFNI (https://afni.nimh.nih.gov) and FreeSurfer v6.0 (https://surfer.nmr.mgh.harvard.edu) for preprocessing, statistical analysis and surface reconstruction. Custom scripts used for coarse-to-fine stimulus generation, threshold-dependent locus identification and atlas-space plotting are available from the corresponding authors on reasonable request.

## Acknowledgements

This work was supported by National Natural Science Foundation of China (32471050 to P.L. and 52293424 to X.Z.), the Fundamental Research Funds for the Central Universities (226-2024-00022, 2025ZFJH01-01 to P.L.), the Non-profit Central Research Institute Fund of Chinese Academy of Medical Sciences (2023-PT310-01 to P.L.), Key R&D Program of Zhejiang Province (2024SSYS0019 to P.L.), Pioneer R&D Program of Zhejiang (2024C03001 to P.L.). Institutional support were provided in part by the National Institutes of Health through the National Institute on Deafness and Other Communication Disorders (R01DC019979) and the National Institute of Mental Health Silvio O. Conte Center (P50MH109429).

## Author contributions

M.Q.: Conceptualization, Methodology, Investigation, Formal analysis, Visualization, Writing – original draft, Supervision.

M.C.: Methodology, Investigation, Formal analysis, Visualization, Writing – review & editing

A.B.: Data interpretation, Writing – review & editing.

X.Z.: RF coil design, MRI acquisition, Writing – review & editing.

P.L.: Resources, Supervision, Writing – review & editing.

A.W.R.: Conceptualization, Methodology, Resources, Supervision, Writing – review & editing.

M.Q. and M.C. contributed equally to this work. M.Q., P.L., and A.W.R. jointly supervised this work and serve as corresponding authors.

## Figures

**Supplementary Fig. 1.**
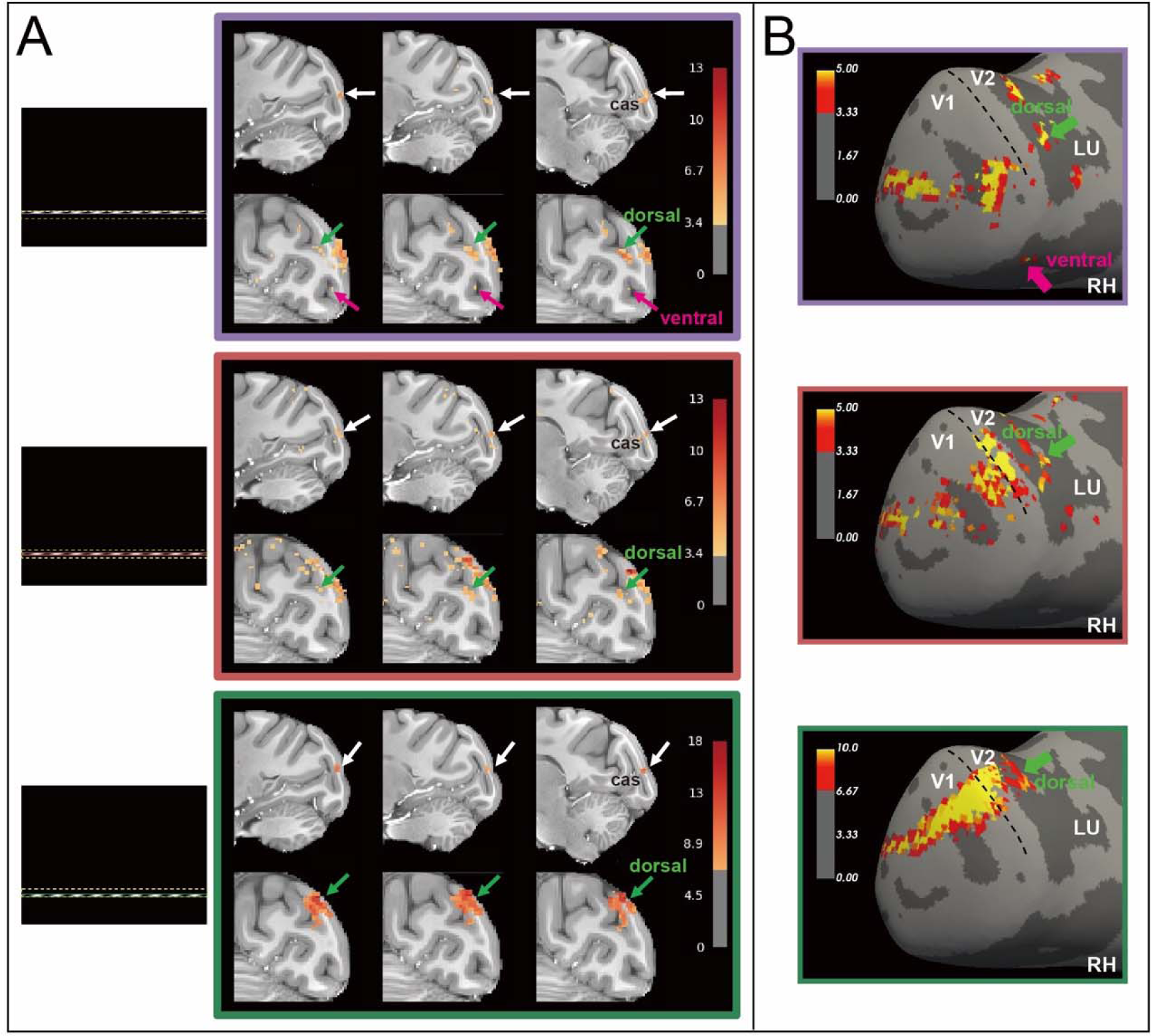
Horizontal meridian mapping with 0.5° wide bar. (A) Horizontal bars were presented at three adjacent vertical positions (upper, middle, and lower), producing a systematic shift of activation between ventral and dorsal portions of early visual cortex. 0.5° wide horizontal bars were presented at adjacent vertical positions (color-coded stimulus positions, shown at left) leads to activations shown in colored frames (purple, red, green). For each condition, sagittal views (upper row) and coronal views (lower row) are shown. White arrows indicate the location of the HM representation in V1, near the calcarine sulcus (labeled cas in the figure). Green arrows indicate dorsal V2/V3 activation and magenta arrows indicate ventral V2/V3 activation. The upper position (purple) achieves the strongest dorsal and ventral activation, while the other two positions are heavily biased towards ventral. (B) Surface views (right hemisphere) for the same three conditions. Green arrows indicate dorsal V2/V3 activation and magenta arrows indicate ventral V2/V3 activation. The upper (purple) condition activated both dorsal and ventral V2/V3 borders and produced the broadest, most spatially continuous response, and largest cortical magnification. LU, lunate sulcus; black dashed line, V1/V2 border. Monkey D. Statistical threshold: p < 0.001.

**Supplementary Fig. 2.**
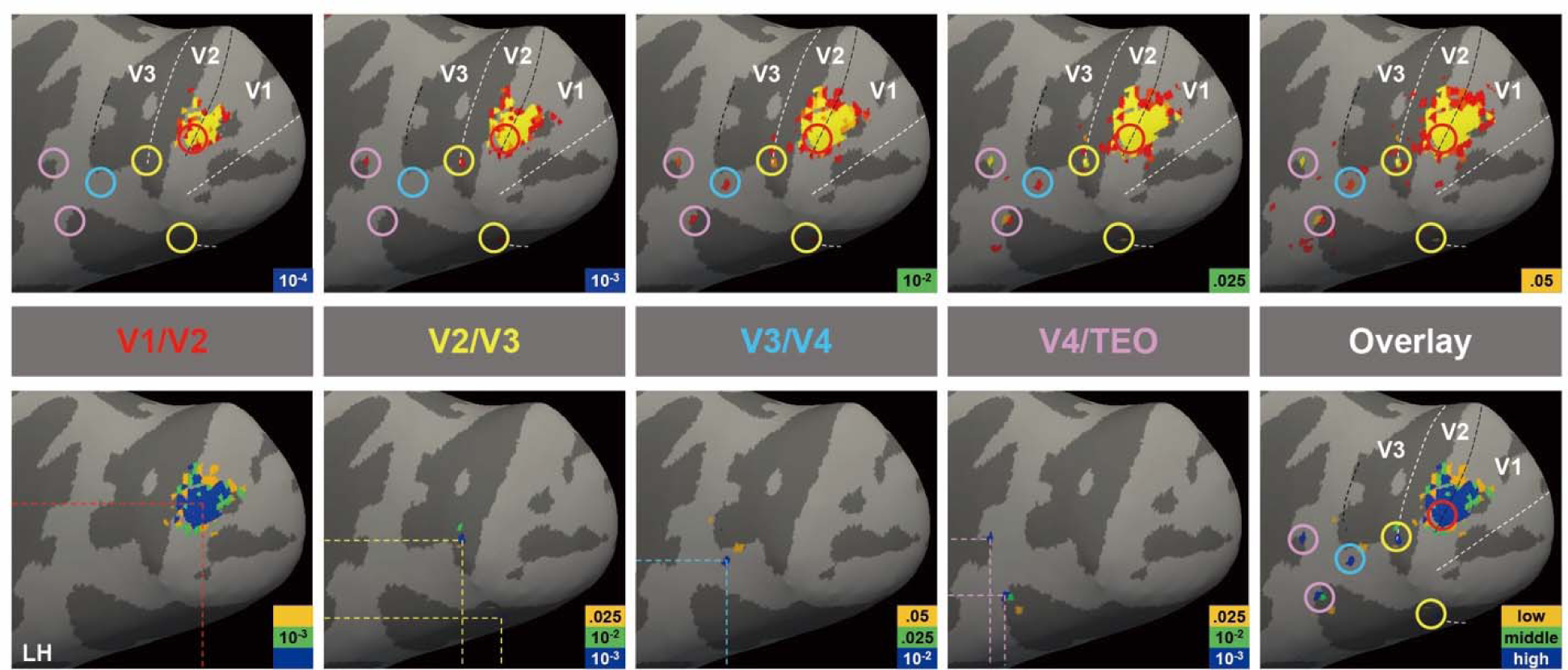
Differential threshold-dependent activation patterns across early visual area borders reveal distributed foveolar loci. Top panels: Cortical surface activation maps evoked by 0.2° foveolar stimulation are shown under two representative statistical thresholds (left: most stringent threshold, p < 10^-4^; middle: intermediate thresholds; right: most liberal threshold, p < 0.05). Colored circles indicate stability of foveolar response loci across multiple threshholds. Bottom panels: Threshold-dependent activation patterns are displayed separately for different areal borders. Left to right: V1/V2 border (red), V2/V3 border (yellow), V3/V4 border (blue), V4/TEO (pink), and their combined overlay (rightmost panel). For each border, three statistical thresholds are shown to illustrate area-specific response strength and stability. Overlay panel: Integration of activation clusters across thresholds and borders reveals multiple foveolar loci distributed along visual cortex boundaries. The presence of consistent activation foci across thresholds supports the stability of foveolar activations and supports distributed foveolar representations as previously found in awake fixating monkeys.

**Supplementary Fig. 3.**
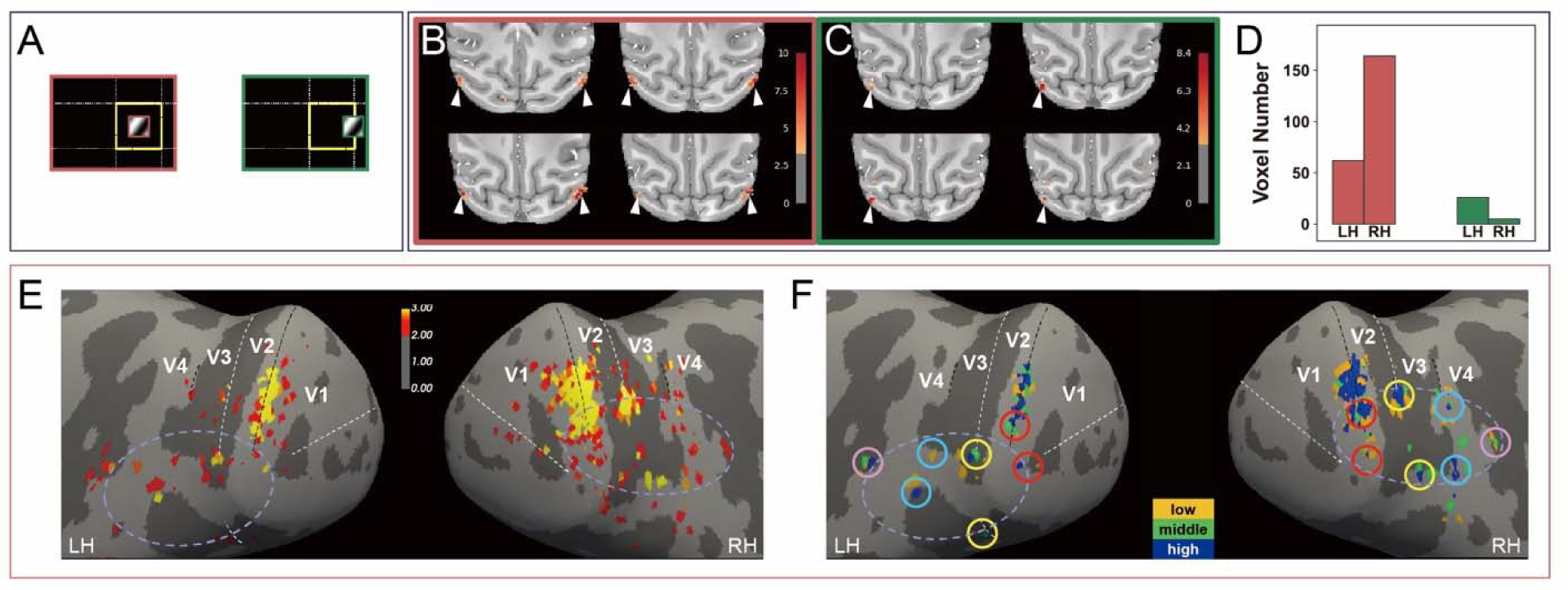
Refining foveolar localization using 0.2° spot stimuli in a second case (Monkey D). (**A**) The 0.2° spot positions presented within the 0.5° region defined by the intersection of the refined VM and HM coordinates; frame color identifies the spot position (purple, red, green) and dotted lines indicate the 0.5° center. (**B, C**) Activation patterns evoked by the centered spot (B, red frame) and by the displaced spot (C, green frame); four horizontal slices through the center of activation are shown for each, and arrowheads indicate activation evoked by the spot. (**D**) Number of activated voxels in the left (LH) and right (RH) hemispheres for each spot position; bar colours correspond to the frame colours in A–C. (**E**) Surface projections of the activation evoked by the centered spot for the left (LH) and right (RH) hemispheres; dashed lines mark the V1/V2, V2/V3 and V3/V4 borders. (**F**) Foveolar loci identified from threshold-dependent activation patterns, with statistical thresholds color-coded and normalized for each cortical area (low: yellow; middle: green; high: blue). Circles mark the foveolar locus for each area and the dashed oval marks the perimeter of the foveolar core. Conventions as in Fig. 5. Statistical threshold for activation maps in B and C: p < 0.001.

**Supplementary Fig. 4.**
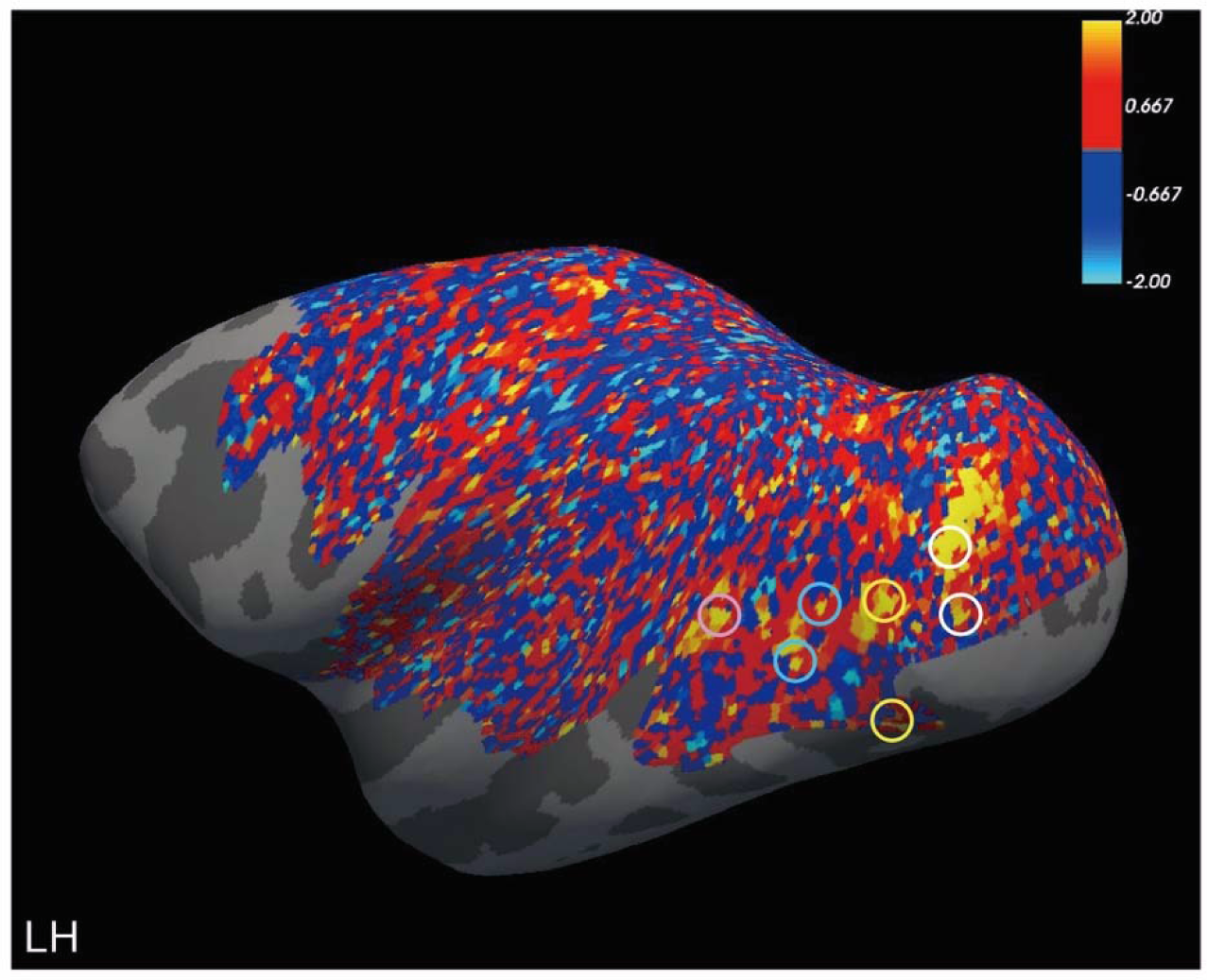
Non-thresholded cortical response map evoked by the central 0.2° spot stimulus in Monkey D. The functional response is projected onto the inflated cortical surface without statistical thresholding to show the complete spatial distribution of stimulus-related signal changes. Colored circles indicate the foveolar loci identified in Fig. 5F. Gray regions denote cortical locations without displayed functional data due to the lack of coil coverage.

